# Combinatorial MAPK states drive opposing gene programmes: a reanalysis of MAP3K-driven transcriptional responses in MCF10A cells

**DOI:** 10.64898/2026.07.10.737700

**Authors:** Mireia Solé, Beatriz Candás-Estébanez, Renata Kelly da Palma, Cristian Pablo Moiola, Noèlia Téllez, Albert Gubern

## Abstract

**Background:** The MAPK signalling network coordinates opposing cell fates — proliferation versus apoptosis — by combining the activities of ERK, JNK, and p38 kinases in patterns that depend on upstream MAP3K identity. Using multiplexed kinase translocation reporter (KTR) biosensors and functional assays in MCF10A mammary epithelial cells, Peterson et al. (Cell Systems, 2023) demonstrated that MAP3Ks co-activating ERK and JNK drive approximately tenfold more cell cycle entry than MAP3Ks activating ERK alone, and that both kinase activities are independently required for this proliferative response. The transcriptional mechanism underlying this quantitative difference was not identified in the original study.

**Results:** We performed a systematic computational re-analysis of the RNA-seq data deposited by Peterson et al. (GEO: GSE213882), encompassing thirteen MAP3K perturbations in doxycycline-induced MCF10A mammary epithelial cells. Based on signalling responses reported by Peterson et al., MAP3Ks were grouped into ERK-only (RAF1, COT) or ERK+JNK co-activating kinases (MLK1, MLK3, ZAK, TAK1, MEKK2, MEKK3). This stratification identified JNK co-activation as the feature most strongly associated with E2F/MYC transcriptional polarity. ERK-only MAP3Ks showed negative enrichment of E2F TARGETS, G2M CHECKPOINT, and MYC TARGETS Hallmark gene sets, whereas ERK+JNK MAP3Ks showed positive enrichment (each MAP3K individually FDR < 0.05 by GSEA). Consistently, inferred activities of E2F1, E2F2, E2F4, and MYC shifted from negative mean ULM t-scores in the ERK group to strongly positive values in the ERK+JNK group (FDR < 0.05 using the CollecTRI regulon). The observed E2F/MYC polarity was robust to alternative transcription factor inference strategies, being reproduced using the DoRothEA A+B regulon and the multivariate model (MLM) framework (Pearson r = 0.77). The signal remained stable under leave-one-out sensitivity analysis and was independently reproduced in MAP3K overexpression signatures from the LINCS L1000 atlas (GEO: GSE92742).

**Conclusions:** ERK activation alone was associated with repression of E2F/MYC transcriptional programmes, whereas JNK co-activation was associated with their activation. These opposing transcriptional states provide a potential explanation for the ∼10-fold differences in cell cycle entry reported by Peterson et al. and supports a model in which pRb/E2F regulation integrates combinatorial MAPK co-activation states beyond the canonical ERK–cyclin D–CDK4/6 axis.

## Introduction

The RAS–ERK signalling axis has long been considered the canonical mitogenic pathway, promoting cell cycle entry via cyclin D accumulation, CDK4/6-mediated phosphorylation of the retinoblastoma protein (pRb), and the consequent release of E2F transcription factors [1,2]. This linear model predicts that ERK activation should be sufficient to drive E2F-dependent gene expression and S-phase commitment. Paradoxically, however, many physiological and pathological stimuli that activate ERK also engage JNK and p38 — the stress-activated MAPK branches — and the net cell fate outcome differs substantially depending on which combination is produced [3,4]. The molecular basis of this combinatorial specificity has remained poorly understood.

The logic connecting upstream MAP3K identity to downstream MAPK combinations and ultimately to cell fate was clarified by Peterson et al. [5], who used multiplexed MAPK kinase translocation reporter (KTR) biosensors to map the MAPK activity outputs across human MAP3Ks. This systematic approach revealed that MAP3Ks segregate into functional groups defined by their MAPK activation signatures: some kinases selectively activate ERK, while others co-activate ERK with JNK, and/or p38. Pharmacological perturbation experiments further demonstrated that p38 co-activation is dispensable for cell cycle entry, establishing ERK and JNK as the critical signalling pair. Critically, ERK+JNK co-activation drives approximately ten-fold more cell cycle entry than ERK activation alone, as measured by EdU incorporation, pRb phosphorylation, and CDK2 activity. Both ERK and JNK activities were independently necessary for this proliferative response, indicating a genuine requirement for combinatorial MAPK signalling rather than simply a difference in ERK amplitude. Despite this key functional insight, the transcriptional mechanism linking JNK co-activation to enhanced proliferation was not defined. The RNA-seq data accompanying this study (GEO: GSE213882) were deposited publicly, providing an opportunity to identify the transcriptional mechanism underlying this quantitative difference in cell fate.

We hypothesised that the ∼10-fold difference in cell cycle entry reflects a qualitative reversal in E2F and MYC transcriptional programme states — the canonical downstream effectors of pRb inactivation — and that this reversal is determined by JNK co-activation status rather than by ERK amplitude alone. Here we present a computational re-analysis of the Peterson et al. RNA-seq dataset integrating Hallmark gene set enrichment, genome-wide fold-change profiling, and transcription factor (TF) activity inference across five independent lines of evidence, including external validation in the LINCS L1000 perturbation atlas and a combined cross-platform consistency analysis. The central finding — that ERK-only MAP3Ks actively repress E2F/MYC programmes while ERK+JNK MAP3Ks activate them — places JNK co-activation as a putative gating mechanism for the E2F transcriptional switch, connecting MAP3K identity to a cell fate decision that precedes and determines cell cycle commitment.

## Results

### MAP3K transcriptional programmes segregate by MAPK co-activation state

We re-processed the RNA-seq count data from Peterson et al. (GEO: GSE213882) using DESeq2. The dataset comprises thirteen MAP3K perturbations plus a shared serum-starved Control in MCF10A cells with doxycycline-inducible MAP3K expression (two biological replicates each). For quality control (QC) and exploratory analysis, counts were variance-stabilised (VST); sample-distance clustering confirmed expected replicate grouping and the dispersion-mean relationship showed well-fitted shrinkage across all MAP3K comparisons (Fig. S1a–b). Differential expression was estimated using DESeq2 Wald statistics, which served as the primary input for gene set enrichment and transcription factor activity analyses.

To obtain an unbiased overview across the full dataset, we performed PCA on VST-normalised counts for all thirteen MAP3K constructs and the Control, using the top 500 most variable genes. PC1 (57% variance explained) was associated with the overall magnitude of the transcriptional response, with Control, NIK, ARAF and BRAF projecting to negative values consistent with minimal transcriptional activation, and MEKK2, MEKK3 and COT projecting to high positive values. The remaining MAP3Ks occupied intermediate positions, reflecting graded responses (Fig. 1a). PC2 (12% variance explained) separated ERK-only MAP3Ks (RAF1, COT) from the ERK+JNK group, with both ERK-only kinases projecting to consistently negative values in contrast to all ERK+JNK MAP3Ks.

**Fig 1.**
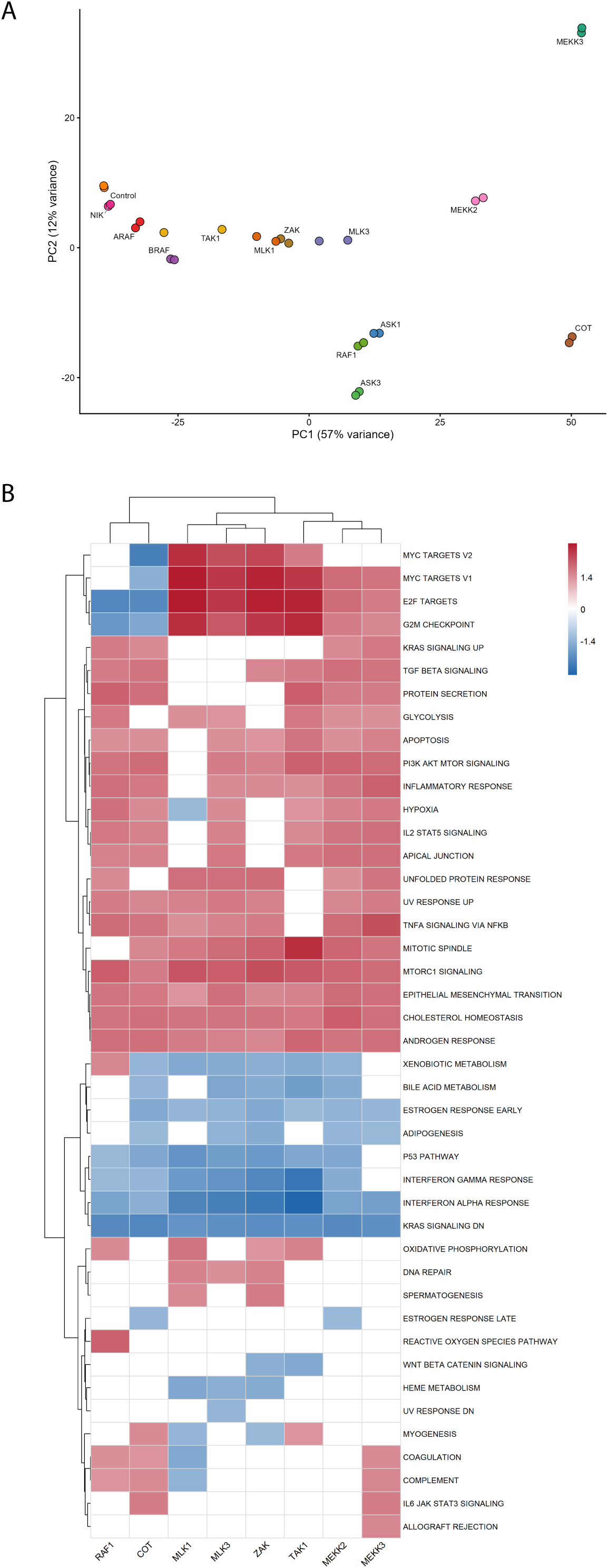
MAP3K transcriptional landscapes segregate by MAPK co-activation state. (a) Principal component analysis (PCA) of VST-normalised RNA-seq counts (top 500 most variable genes by row variance). All thirteen MAP3K perturbations and serum-starved Control are shown (n = 2 biological replicates per condition). Each point represents one sample; colour indicates MAP3K identity. Percentage of variance explained by each PC is shown on the axis labels. (b) Hallmark gene set enrichment heatmap. NES from GSEA (fgsea; Hallmark v7.4) for all 50 Hallmark pathways across the eight classified MAP3Ks, ranked by DESeq2 Wald t-statistic. White: FDR ≥0.05. Columns and rows clustered by Ward’s method.

To test whether MAPK co-activation state structures transcriptional output, MAP3Ks were classified into two groups based on KTR activity profiles reported by Peterson et al. [5] (Fig. S2). Five MAP3Ks were excluded on pre-specified, a priori grounds. ARAF and NIK were excluded due to absence of detectable ERK or JNK activity. ASK1 and ASK3 were excluded despite measurable p38 activity, as neither kinase produces detectable ERK or JNK output, and p38 co-activation alone is dispensable for MAP3K-driven cell cycle entry in this system [5]. BRAF was excluded on two independent grounds: its transcriptional profile clusters with kinase-inactive MAP3Ks in the PCA (Fig. 1a), and Peterson et al. demonstrated that its apparent ERK activity is non-cell-autonomous, dependent on paracrine EGFR signalling rather than direct MAP3K output [5]. The remaining eight MAP3Ks were classified into two groups: an ERK-only group comprising MAP3Ks with substantial ERK activity and negligible JNK co-activation (RAF1, COT; n = 2), and an ERK+JNK group comprising MAP3Ks that co-activate both ERK and JNK (MLK1, MLK3, ZAK, TAK1, MEKK2, MEKK3; n = 6).

To characterise transcriptional programmes associated with MAPK co-activation state, we performed GSEA with the full Hallmark collection, using DESeq2 Wald statistics as input for each of the eight classified MAP3Ks. Among all 50 Hallmark gene sets, E2F TARGETS, G2M CHECKPOINT and MYC TARGETS showed the most consistent directional separation between signalling groups (Fig. 1b). ERK+JNK MAP3Ks consistently exhibited positive enrichment across all three gene sets (individual FDR < 0.05), whereas ERK-only MAP3Ks showed negative enrichment for E2F TARGETS and G2M CHECKPOINT (individual FDR < 0.05). The direction of enrichment was consistent across all MAP3Ks within each signalling class. In contrast, several Hallmark pathways — including MTORC1 SIGNALING — showed consistently positive enrichment across all eight MAP3Ks regardless of JNK co-activation status (Fig. 1b), indicating that MAP3K overexpression elicited substantial transcriptional responses in both groups and suggesting that the E2F/MYC polarity reflects a specific feature of MAPK co-activation state rather than a global difference in transcriptional output.

### JNK co-activation is associated with E2F/MYC transcriptional polarity

To quantify the transcriptional polarity suggested by the GSEA analysis, we compared NES values for three a priori-selected Hallmark gene sets between the ERK group (n = 2: RAF1, COT) and the ERK+JNK group (n = 6: MLK1, MLK3, ZAK, TAK1, MEKK2, MEKK3) (Fig. 2a). E2F TARGETS, G2M CHECKPOINT, and MYC TARGETS were selected before group-level statistical testing based on their functional relevance to the cell-cycle entry phenotype reported by Peterson et al.[5]. G2M CHECKPOINT was included because it shares a substantial fraction of its leading-edge genes with E2F TARGETS and captures the broader S/G2 transcriptional programme driven by E2F1/E2F4, thereby providing an orthogonal confirmation of the observed polarity.

**Fig 2.**
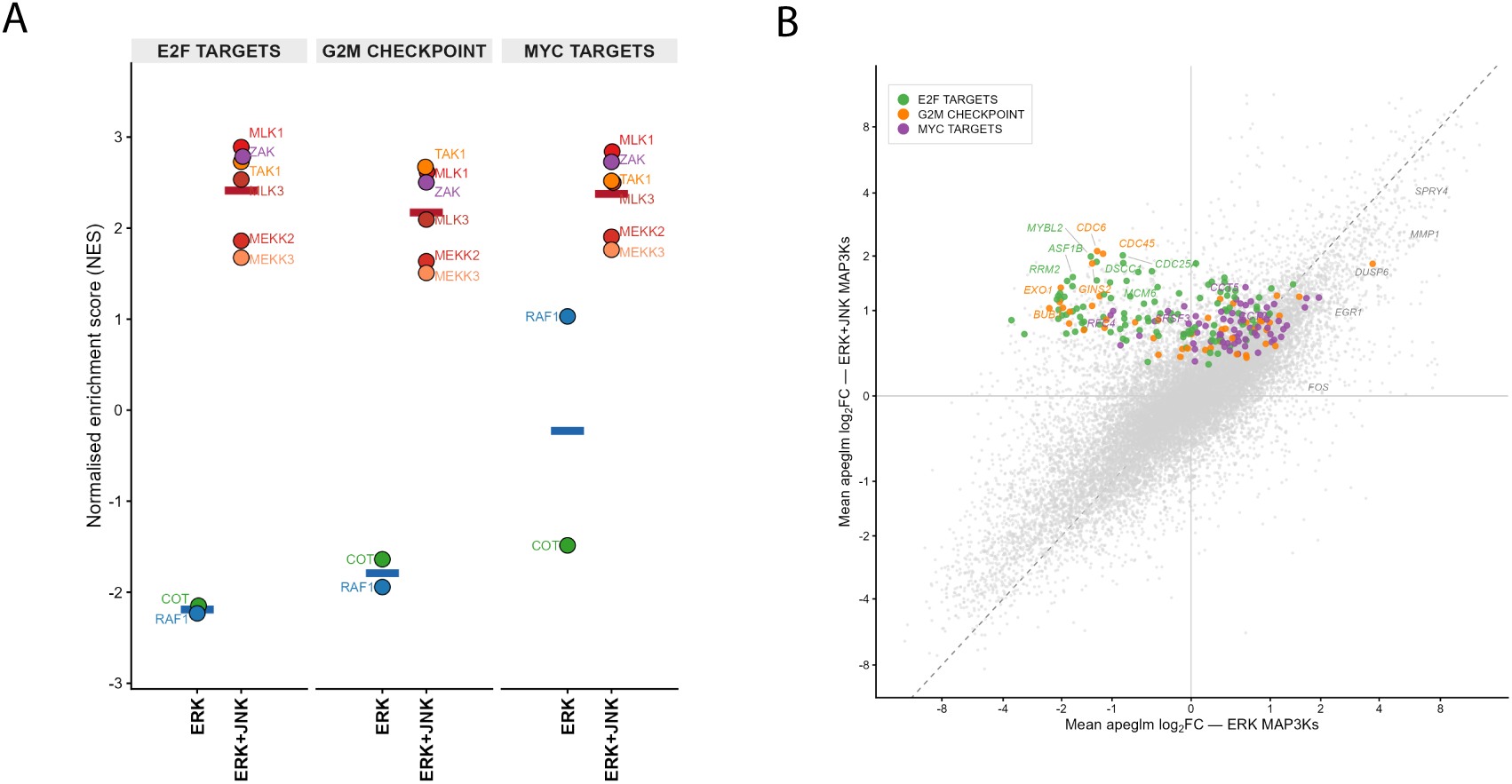
JNK co-activation determines the polarity of E2F/MYC transcriptional programmes. (a) Beeswarm plot of Hallmark NES values by group. Each point represents one MAP3K (ERK group: RAF1, COT, n = 2; ERK+JNK group: MLK1, MLK3, ZAK, TAK1, MEKK2, MEKK3, n = 6). Crossbars indicate group means. (b) Genome-wide fold-change scatter plot. Each point represents one expressed gene (n = 21,124). X-axis: mean apeglm-shrunk log₂ fold-change across ERK-only MAP3Ks vs Control. Y-axis: mean log₂ fold-change across ERK+JNK MAP3Ks vs Control. Both axes on asinh scale. Dashed diagonal: y = x. Coloured points: genes in the Hallmark leading edges of E2F TARGETS (green), G2M CHECKPOINT (orange), or MYC TARGETS (purple), present in ERK+JNK MAP3Ks. Grey italic labels: canonical ERK immediate-early genes (EGR1, FOS, DUSP6, SPRY4, MMP1).

A striking directional separation was observed between signalling groups. E2F TARGETS showed uniformly negative enrichment among ERK-only MAP3Ks (mean NES = −2.19) and uniformly positive enrichment among ERK+JNK MAP3Ks (mean NES = +2.41), corresponding to a difference of Δ NES = +4.60. Similarly, G2M CHECKPOINT displayed negative enrichment in the ERK group (mean NES = −1.79) and positive enrichment in the ERK+JNK group (mean NES = +2.17, Δ NES = +3.96). For both pathways, no overlap in NES values was observed between signalling classes. MYC TARGETS showed the same overall directional trend (ERK mean NES = −0.23, ERK+JNK mean NES = +2.38, ΔNES = +2.60), although the response was more heterogeneous within the ERK-only group, with RAF1 exhibiting a weak positive enrichment and COT a strong negative enrichment.

The genome-wide dimension of this polarity is illustrated by a scatter plot comparing mean apeglm-shrunk log₂ fold-change across ERK+JNK MAP3Ks (y-axis) versus ERK MAP3Ks (x-axis) for all 21,124 expressed genes (Fig. 2b, asinh scale). Most genes distributed along the diagonal, indicating a substantial shared global activation response across both signalling states. In contrast, leading-edge genes from E2F TARGETS, G2M CHECKPOINT, and MYC TARGETS clustered preferentially in the upper-left quadrant, corresponding to induction in ERK+JNK conditions but repression or minimal activation in the ERK-only group. This pattern demonstrates that the observed E2F/MYC reflects coordinated regulation across numerous target genes rather than the behaviour of a small subset of genes highly influential genes. Notably, canonical ERK-regulated genes, including transcriptional feedback targets (DUSP6, SPRY4) and AP-1 components (FOS, EGR1, MMP1) localised along the diagonal or within the positive x-axis region, indicating robust ERK pathway activation in both RAF1 and COT conditions. Together, these observations suggest that the repression of E2F-associated programmes in the ERK-only group cannot be attributed to absent or defective canonical ERK signalling, but instead reflects a specific transcriptional feature associated with MAPK-coactivation state.

### E2F1, E2F2, E2F4 and MYC transcriptional activity is associated with MAPK co-activation state

GSEA identifies programme-level enrichment but does not resolve which specific transcription factors drive the observed polarity, nor whether these activity patterns are accompanied by corresponding changes in TF transcript abundance and activity. To address this, we inferred per-MAP3K TF activity using the Univariate Linear Model (ULM) [6] with the CollecTRI signed TF-target regulon [7]. Input was the DESeq2 Wald t-statistic per gene, which provides gene-level signal-to-noise weighting appropriate for low-replicate designs. ULM t-scores reflect the linear association between each gene’s differential expression rank and its signed weight in the CollecTRI regulon. Four focus TFs were pre-specified from the Hallmark results in Fig. 2: E2F1, E2F2, and E2F4, representing the E2F TARGETS and G2M CHECKPOINT programmes, and MYC, representing the MYC TARGETS.

Consistent with the pathway-level analyses, all four TFs exhibited a clear polarity according to MAPK co-activation state (Fig. 3a). E2F1 activity was negative in both ERK-only MAP3Ks (RAF1: t = −5.65, FDR < 0.001; COT: t = −3.74, FDR < 0.01) but positive in all ERK+JNK MAP3Ks (range +4.96 to +11.58, all FDR < 0.001), resulting in group means of −4.70 and +7.91, respectively. Similar patterns were observed for E2F4 (group means: −4.16 versus +6.75; all ERK+JNK individually FDR < 0.05) and E2F2 (−3.60 versus +4.65; all ERK+JNK individually FDR < 0.05 except TAK1, FDR = 0.09). MYC showed the same overall directional pattern (group means: −1.80 versus +7.37; all ERK+JNK individually FDR < 0.001), although RAF1 displayed near-neutral MYC activity (t = +0.30, FDR = 0.76) whereas COT showed significant repression (t = −3.89, FDR < 0.01); this divergence is consistent with the NES values shown in Fig. 2a, where RAF1 shows near-neutral MYC TARGETS enrichment compared with significant repression by COT.

**Fig 3.**
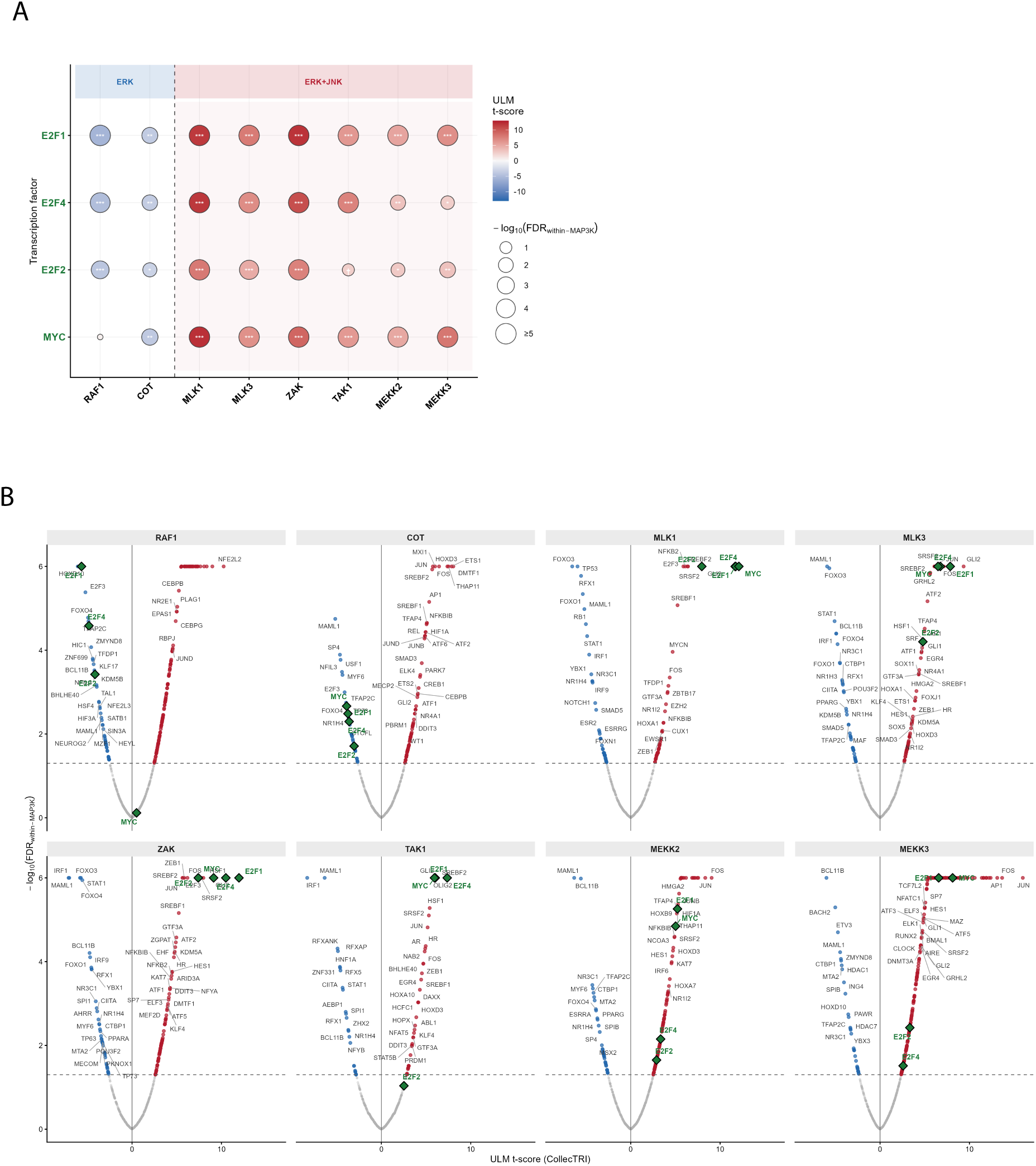
E2F and MYC transcription factor activity switches are gated by JNK co-activation. (a) Dot plot of ULM t-scores for pre-specified focus TFs. TF activity inferred using ULM (decoupleR) with the CollecTRI signed TF-target regulon. Input: DESeq2 Wald t-statistics per gene per MAP3K. Four focus TFs pre-specified from Fig. 2: E2F1, E2F4, E2F2 (E2F TARGETS/G2M CHECKPOINT), and MYC (MYC TARGETS). Fill colour: ULM t-score (blue = repression; red = activation). Point size: −log₁₀(within-MAP3K BH FDR), capped at 5. Dashed vertical line separates ERK (left, blue header) from ERK+JNK (right, red header) groups. † FDR < 0.10; * FDR < 0.05; ** FDR < 0.01; *** FDR < 0.001. (b) Volcano plot of ULM t-scores across all CollecTRI TFs. Each point represents one TF (n > 700 per MAP3K). X-axis: ULM t-score. Y-axis: −log₁₀(within-MAP3K BH FDR), capped at 5. Panels faceted by MAP3K; column order matches (a). Focus TFs labelled and coloured. Dashed horizontal line: FDR = 0.05.

Notably, these large ULM t-score differences stand in sharp contrast to the variable and directionally inconsistent changes in E2F1, E2F4, E2F2, and MYC mRNA abundance across MAP3Ks (Supplementary Fig. S3a). This dissociation suggests that transcriptional upregulation of these TF genes is not the primary driver of programme activation and is instead consistent with regulatory mechanisms operating downstream transcription.

The transcription-wide context of these findings is illustrated by volcano plots showing inferred activity for all CollecTRI transcription factors across each MAP3K perturbation (Fig. 3b). In an unbiased analysis of more than 700 TFs, E2F1, E2F4, E2F2, and MYC consistently emerge among the most positively inferred regulators in ERK+JNK MAP3Ks and among the most negatively inferred regulators in ERK MAP3Ks. This observation indicates that the E2F/MYC polarity is not an artefact of TF pre-selection but represents one of the dominant regulatory features distinguishing the two signalling states.

Several analyses supported the robustness of this finding. First, leave-one-out sensitivity analysis showed that no single MAP3K drives the group-level conclusion: the maximum change in group mean delta across all focus TFs was 0.91 t-score units (E2F4, removing MLK1), well within the 95% confidence interval of the full-dataset estimate (Supplementary Fig. S3b). Second, repeating TF activity inference with the DoRothEA A+B regulon [8] and MLM estimator yielded highly concordant results (Pearson r = 0.77, p < 0.001, n = 32 TF × MAP3K pairs; Supplementary Fig. S3c). Together, these findings indicate that activation of E2F-driven transcriptional programmes is a consistent feature of ERK+JNK signalling, whereas ERK-only signalling is associated with repression of these programmes.

### External validation in the LINCS L1000 perturbation atlas supports E2F/MYC polarity

To assess the generalisability of the E2F/MYC polarity identified in the Peterson dataset, we analysed MAP3K overexpression signatures from the LINCS L1000 Phase I atlas (GEO: GSE92742) [9]. Four MAP3Ks were represented as overexpression perturbagens: RAF1 (ERK group; consensus across n = 10 cell lines), COT (ERK group; n = 10 cell lines), MEKK2 (ERK+JNK group; n = 1 cell line), and MLK1 (ERK+JNK group; n = 1 cell line). Consensus expression profiles were generated by averaging COMPZ scores across available cell lines for each perturbagen.

GSEA reproduced the directional pattern observed in the primary dataset (Fig. 4). Both ERK-only MAP3Ks showed significant negative enrichment of E2F TARGETS and G2M CHECKPOINT (RAF1: NES = −2.88 and −2.41; COT: NES = −1.89 and −1.70, respectively; all FDR < 0.001). RAF1 additionally showed significant negative enrichment of MYC TARGETS (NES = −1.71, FDR < 0.001), whereas MYC TARGETS was not significant for COT in the LINCS landmark gene space (NES = +0.97, FDR = 0.64). In contrast, MEKK2 exhibited strong positive enrichment for all three gene sets (E2F TARGETS NES = +2.74, G2M CHECKPOINT NES = +2.81, MYC TARGETS NES = +3.59; all FDR < 0.001). MLK1 showed positive enrichment of E2F TARGETS (NES = +1.42, FDR < 0.05), although G2M CHECKPOINT and MYC TARGETS did not reach FDR < 0.05 for this single-cell-line consensus.

**Fig 4.**
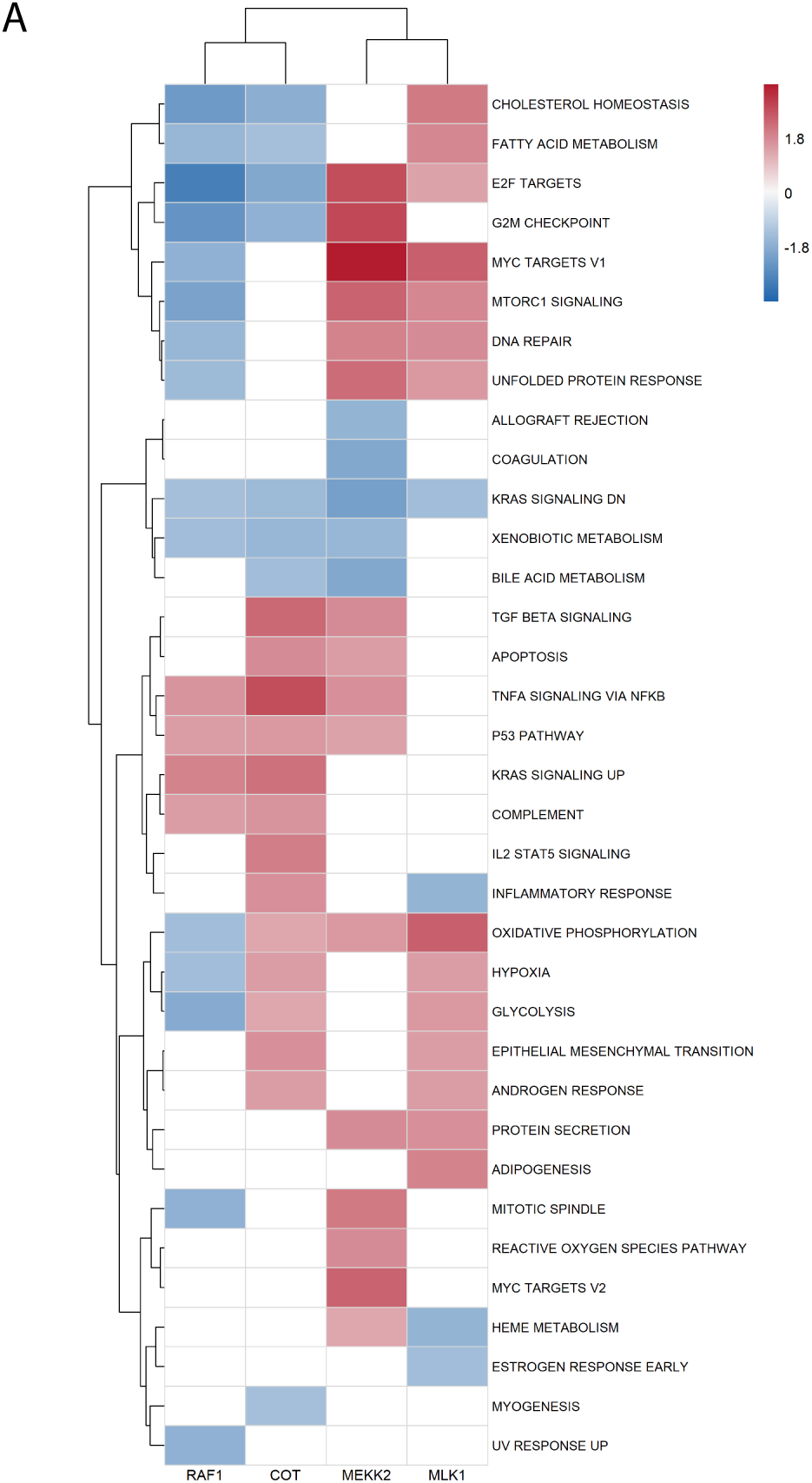
External validation of E2F/MYC polarity in the LINCS L1000 perturbation atlas. GSEA Hallmark heatmap — LINCS L1000. MAP3K overexpression signatures from LINCS L1000 Phase I (GEO: GSE92742; RAF1, COT: consensus across n = 10 cell lines each; MEKK2, MLK1: n = 1 cell line each). NES encoding as in Fig. 1b; white = FDR ≥0.05.

TF activity inference also yielded a consistent directional pattern (Supplementary Fig. S4a). Among ERK+JNK MAP3Ks, MEKK2 showed strong positive activity for all four focus TFs (E2F1: t = +4.45, FDR < 0.001; E2F4: t = +5.79, FDR < 0.001; E2F2: t = +4.06, FDR < 0.001; MYC: t = +6.76, FDR < 0.001). MLK1 provided partial support, with significant activation of MYC (t = +5.28, FDR < 0.001) and E2F4 (t = +2.63, FDR = 0.017), whereas E2F1 and E2F2 did not reach FDR < 0.05 for this single-cell-line consensus. Conversely, RAF1 showed negative activity for E2F1 (t = −4.90, FDR < 0.001), E2F4 (t = −3.44, FDR < 0.01), E2F2 (t = −2.80, FDR < 0.01) and a negative trend for MYC (t = −1.93, FDR = 0.053). COT showed significant repression of E2F4 (t = −2.52, FDR < 0.05); remaining focus TFs were near-neutral and non-significant. In the unbiased volcano across all CollecTRI TFs (Supplementary Fig. S4b), E2F1, E2F4, E2F2 and MYC emerge at the extreme positive end for MEKK2 and at predominantly negative positions for RAF1, without prior selection, independently reproducing the bidirectional pattern from Fig. 3b.

We acknowledge two limitations of this external validation. First, LINCS landmark space (∼978 genes) covers a fraction of the full transcriptome; CollecTRI regulon coverage was confirmed above the minimum ULM threshold for all focus TFs (see Methods). Second, the ERK+JNK arm rests on a single cell line per MAP3K, preventing cross-cell-line consistency assessment for this group, in contrast to the ten-cell-line ERK consensus which provides robust cross-cell-line evidence for E2F repression by ERK-only signals. Despite this asymmetry, E2F1, E2F2, E2F4 and MYC emerged among the highest-ranked activated regulators in MEKK2 and among the most negatively ranked regulators in RAF1 in an unbiased scan of over 700 transcription factors. Consequently, the LINCS analysis should be viewed as supportive rather than definitive validation, while nevertheless providing evidence that the observed E2F/MYC polarity extends across independent datasets and experimental systems.

### Cross-platform consistency supports the robustness of the E2F/MYC polarity

To assess cross-platform consistency, we pooled MAP3K observations from the Peterson RNA-seq and LINCS L1000 datasets into a single combined analysis, treating each MAP3K × dataset combination as one observation. This approach treats both platforms symmetrically, without privileging either dataset, while acknowledging that LINCS landmark gene space (∼978 genes) and the Peterson full transcriptome (∼20,000 genes) are not directly comparable in scale; the purpose of this analysis is to confirm directional consistency across platforms, not to increase precision. ERK group: n = 4 (RAF1 and COT, one observation per platform); ERK+JNK group: n = 8 (six Peterson MAP3Ks plus MEKK2 and MLK1 from LINCS). We note that RAF1 and COT each contribute one observation per dataset and are therefore not fully independent; results should be interpreted as evidence of directional consistency across platforms rather than as independent replications. Two-sided exact Wilcoxon rank-sum tests with Benjamini-Hochberg FDR were applied to GSEA NES values and ULM t-scores independently.

Across both datasets, ERK-only MAP3Ks showed negative enrichment of E2F TARGETS and G2M CHECKPOINT programmes and negative inferred activity of E2F family transcription factors, whereas ERK+JNK MAP3Ks showed positive enrichment and positive activity scores. MYC showed the same overall directional pattern, although with greater variability among individual MAP3Ks. All three proliferation Hallmark gene sets remained significant in the combined analysis (Fig. 5A): E2F TARGETS (mean ERK NES = −2.29, mean ERK+JNK NES = +2.33; FDR < 0.01), G2M CHECKPOINT (−1.92; +2.10; FDR < 0.01), MYC TARGETS (−0.30; +2.54; FDR < 0.01). At the TF level, all four focus TFs remained significant (Fig. 5B): E2F1 (mean ERK t = −3.46, ERK+JNK t = +6.50; FDR < 0.01), E2F4 (−3.57; +6.11; FDR < 0.01), E2F2 (−2.57; +3.99; FDR < 0.01), MYC (−1.17; +7.03; FDR < 0.01).

**Fig 5.**
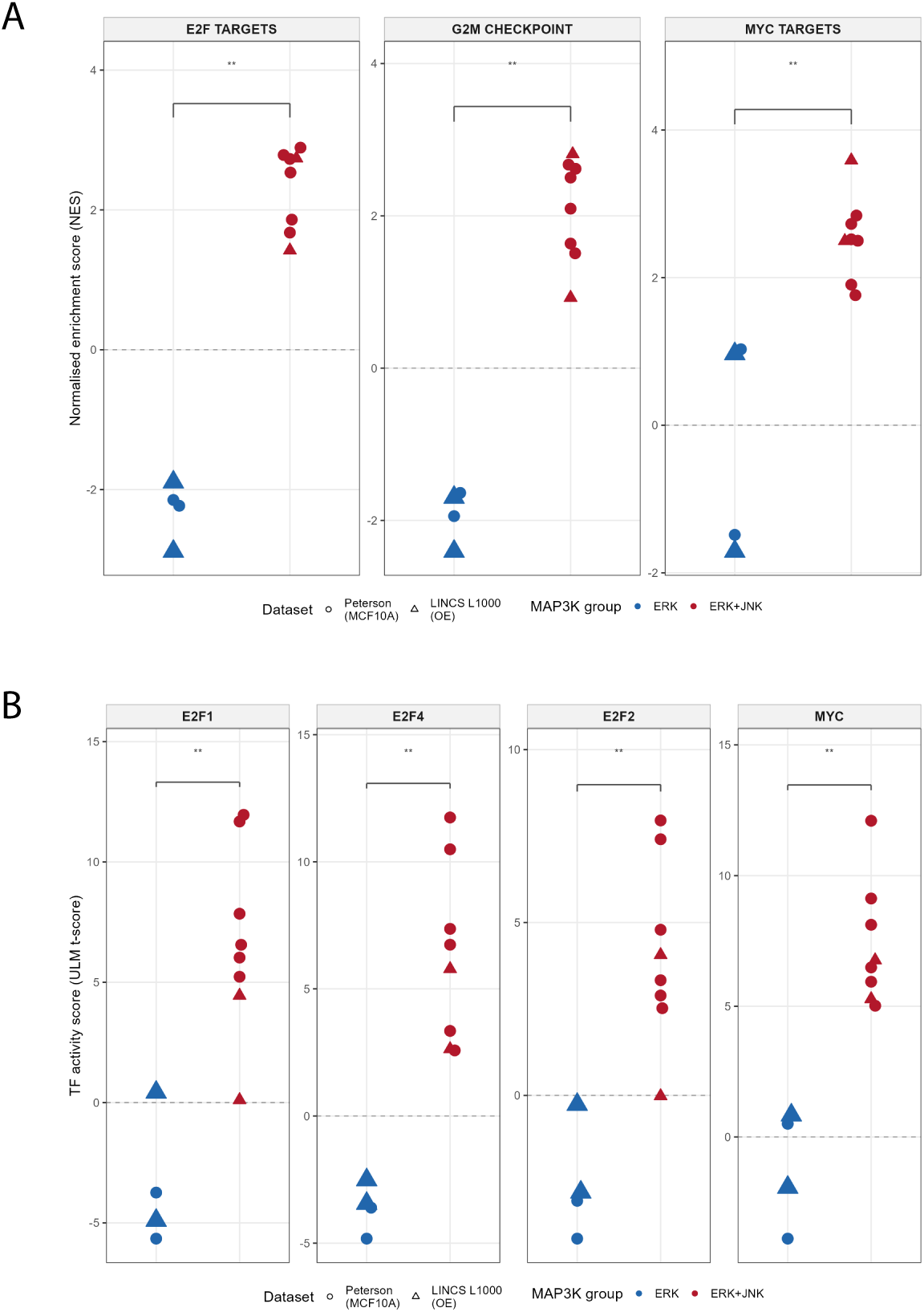
Cross-dataset consistency confirms ERK repression and ERK+JNK activation across experimental platforms. (A) Beeswarm plot of GSEA Hallmark NES values — combined Peterson+LINCS dataset. Three pre-specified proliferation gene sets (ERK n = 4; ERK+JNK n = 8). Circles: Peterson MCF10A RNA-seq; triangles: LINCS L1000 overexpression signatures. ** FDR < 0.01 (exact Wilcoxon rank-sum test, BH correction). (B) Beeswarm plot of ULM t-scores — combined Peterson+LINCS dataset. Four focus TFs. Layout and encoding as in (A). ** FDR < 0.01.

Collectively, the preservation of the E2F/MYC polarity across distinct datasets, analytical frameworks, and transcriptional profiling platforms supports the robustness of the association between ERK-only signalling and negative E2F/MYC enrichment, and ERK+JNK signalling and positive E2F/MYC enrichment, independently of the experimental platform.

## Discussion

This re-analysis of the Peterson et al. RNA-seq dataset reveals a central and previously uncharacterised feature of MAP3K signalling: MAP3Ks that activate ERK alone are associated with repression of E2F/MYC transcriptional programmes, whereas MAP3Ks that co-activate ERK and JNK are associated with activation of these programmes. This is not merely a quantitative difference in transcriptional output — it is a reversal in the direction of regulation. E2F TARGETS and G2M CHECKPOINT exhibited complete directional separation between signalling classes, with all ERK-only MAP3Ks showing negative enrichment and all ERK+JNK MAP3Ks showing positive enrichment; a similar pattern was observed for MYC-associated programmes and for inferred activity of E2F1, E2F2, E2F4 and MYC. The consistency of this polarity across pathway-level enrichment analysis, transcription factor activity inference, alternative regulon resources, and independent validation in the LINCS L1000 perturbation atlas identifies E2F/MYC regulation as a prominent transcriptome-wide feature associated with MAPK co-activation state, and provides a potential mechanistic explanation for the ∼10-fold differences in cell cycle entry reported by Peterson et al. These findings suggest that the pRb/E2F axis in this context operates as a combinatorial integrator of MAPK network state — extending the canonical ERK–cyclin D–CDK4/6–E2F model [1,10] to incorporate JNK co-activation as a candidate gating signal.

A key mechanistic question raised by these findings is why ERK activation alone fails to drive — and is associated with repression of — E2F/MYC programmes. In quiescent MCF10A cells, constitutive MAP3K overexpression represents sustained, non-receptor-coupled kinase activity; sustained ERK signalling is well established to engage strong negative feedback through ERK-driven induction of DUSPs and Sprouty family proteins that attenuate upstream pathway activity [11]. This negative feedback topology, combined with the bistable, switch-like nature of the pRb/E2F circuit [12,13], may explain why partial or feedback-dampened ERK signalling is insufficient to hyperphosphorylate pRb and cross the bistable threshold for E2F release. The observation that RAF1 and COT — two structurally distinct MAP3Ks with different effector specificities — display similar E2F repression across both the Peterson and LINCS datasets is consistent with the possibility that this behaviour reflects a general property of sustained ERK signalling rather than an idiosyncrasy of any single kinase or cellular context.

The nature of this putative post-translational switch is further illuminated by the dissociation between TF activity scores and TF mRNA levels, which argues against transcriptional upregulation of E2F family members as the primary mechanism. This dissociation suggests that JNK co-activation gates E2F programme activity through mechanisms acting downstream of TF transcription. Potential mechanisms include pRb phosphorylation-mediated E2F release, MYC protein stabilisation, cyclin E/CDK2 activation, or other post-transcriptional and post-translational regulatory processes. The most parsimonious model consistent with our data is that ERK-driven cyclin D/CDK4/6 produces partial phosphorylation of pRb that is insufficient alone to trigger full E2F release in quiescent MCF10A cells. In this framework, additional JNK-associated inputs cyclin E/CDK2 activation [14] and chromatin remodelling at E2F target loci — may collectively contribute to crossing the bistable threshold and stabilising the transcriptional switch. An additional possibility is direct JNK-mediated phosphorylation of pRb or components of the pRb–E2F complex, analogous to stress-kinase regulation of pRb described for p38 [15]. These possibilities should be considered hypotheses generated by the present analysis and require direct experimental testing.

Several limitations should be acknowledged. The ERK-only group comprised only two MAP3Ks, and each perturbation was represented by two biological replicates in the original RNA-seq dataset. The LINCS validation dataset provided limited representation of ERK+JNK MAP3Ks, preventing robust assessment of cross-cell-line reproducibility for this signalling class. In addition, both pathway enrichment and TF activity analyses rely on computational inference and therefore cannot establish causality. Nevertheless, the convergence of evidence across independent datasets, regulon resources, inference methods, kinase subsets and sensitivity analyses supports the robustness of the observed E2F/MYC polarity.

Beyond the primary E2F/MYC axis, the divergence of MYC regulation between the two ERK-only MAP3Ks — RAF1 showing near-neutral activity versus COT’s significant repression — is worth noting as a secondary observation. COT has well-characterised roles in TNFα-induced ERK activation and inflammatory signalling [16] and may engage effector contexts that contribute to MYC target repression beyond canonical ERK output, suggesting that MYC regulation may be more context-dependent than E2F regulation downstream of MAP3K signalling.

In cancer biology, the E2F-gating association described here may help explain why RAF1 and COT mutations, which activate ERK without JNK co-activation, require additional alterations to achieve full cell cycle deregulation [17]; conversely, MLK1, which carries somatic loss-of-function mutations in metastatic melanoma [18], may suppress proliferation partly through attenuation of ERK+JNK-driven E2F activation. These associations are correlative, but they generate specific, testable predictions: JNK inhibition in ERK+JNK-co-activating MAP3K contexts should be sufficient to collapse E2F programme activation, and somatic MLK1 loss-of-function should attenuate ERK+JNK-driven E2F/MYC activity in melanoma contexts where JNK co-activation is otherwise intact.

This study demonstrates that publicly available functional genomics data, when re-analysed with systematic regulatory network inference approaches, can generate testable mechanistic hypotheses. The analytical framework — GSEA for hypothesis generation, ULM for TF-level inference, cross-dataset validation in LINCS L1000, and combined cross-platform consistency analysis — is broadly applicable to other perturbation RNA-seq datasets involving combinatorial signalling inputs.

## Materials and Methods

### Source data and experimental design

RNA-seq data were obtained from GEO (accession GSE213882; Peterson et al. [5]). Per-sample transcript-level count files were provided for MCF10A human mammary epithelial cells with doxycycline-inducible TRE3G::MAP3K expression (4CTet system), serum-starved for 24 hours with MAP3K induced by doxycycline (2 μg/mL) for 18 hours. The design comprises thirteen MAP3K conditions plus a shared serum-starved Control (two biological replicates each). For external validation, Level-5 COMPZ moderated z-scores from LINCS L1000 Phase I (GEO: GSE92742) [9] were used as described below.

### MAP3K group assignment

MAP3Ks were assigned to ERK-only or ERK+JNK groups based on KTR activity thresholds defined a priori by Peterson et al. [5]: ERK+JNK, KTR ERK ≥0.5 AND KTR JNK ≥0.5 (MLK1, MLK3, ZAK, TAK1, MEKK2, MEKK3); ERK-only, KTR ERK ≥0.5 AND KTR JNK < 0.5 (RAF1, COT). Excluded MAP3Ks and rationale are detailed in Results. MEKK2 and MEKK3 were retained despite sub-maximal p38 (KTR p38 = 0.7 and 0.4) because p38 inhibition does not impair ERK+JNK-driven proliferation [5].

### Data processing and quality control

Transcript counts were summed to gene level by HGNC symbol. A single DESeq2 [19] object was built for variance-stabilising transformation (VST) used exclusively for QC and PCA. QC included sample-distance heatmaps and dispersion-mean relationship plots. PCA was computed on the 500 most variable genes by row variance of VST expression.

### Differential expression and fold-change shrinkage

For each MAP3K, a separate DESeq2 model was fitted on the subset of samples including that MAP3K and the shared Control (design: ∼ group; reference level: Control). Genes with fewer than 5 counts in at least 2 samples were excluded prior to fitting. Differential expression was estimated using the Wald test; apeglm shrinkage was applied for visualisation and genome-wide scatter [20]. Unshrunken Wald t-statistics (LFC/lfcSE) were used as input to ULM and GSEA. Within-comparison Benjamini-Hochberg FDR correction was applied.

### Gene Set Enrichment Analysis

GSEA was performed using fgsea [21] (eps = 0; minSize = 15; maxSize = 500) with the MSigDB Hallmark collection v7.4 [22], downloaded via msigdbr. Genes were ranked by DESeq2 Wald t-statistic. All 50 Hallmark gene sets were tested; no pre-selection was applied for the overview heatmap. Three gene sets were pre-specified for the group-level test: E2F TARGETS, G2M CHECKPOINT, and MYC TARGETS V1. MYC TARGETS V2 was excluded from the primary test due to near-redundancy with V1 (NES r = 0.984 across MAP3Ks).

### TF activity inference

TF activity was estimated using ULM [6] from the decoupleR package with the CollecTRI regulon [7] (downloaded via OmnipathR [23]; split_complexes = FALSE, retaining protein complexes as single regulatory units to preserve the biological context of cooperative TF binding; activating and repressing weights retained). Input: DESeq2 Wald t-statistic. Minimum regulon size: 5 targets. FDR: Benjamini-Hochberg within each MAP3K. Four focus TFs were pre-specified from Hallmark results: E2F1, E2F2, E2F4, and MYC. Method concordance was assessed by repeating TF activity inference with an independent regulon and estimator: DoRothEA A+B [8], which derives TF-target interactions from curated literature and ChIP-seq evidence ranked by confidence (levels A and B retained), combined with the Multivariate Linear Model (MLM) estimator from decoupleR, which accounts for co-regulation among targets within the same regulon. Concordance between ULM+CollecTRI and MLM+DoRothEA was evaluated by Pearson correlation across all focus TF × MAP3K pairs.

### Leave-one-out sensitivity analysis

Each ERK+JNK MAP3K was removed in turn (n = 6 iterations); group mean t-scores and the ERK+JNK versus ERK group difference (Δ) were recomputed for each focus transcription factor. Changes in the resulting group differences were used to evaluate the influence of individual MAP3Ks on the overall conclusion.

### LINCS L1000 external validation

Level-5 COMPZ z-scores (GEO: GSE92742) were read using cmapR [24]. Overexpression signatures for RAF1, COT, MEKK2, and MLK1 were selected from sig_info metadata. Z-scores were averaged per MAP3K across cell lines (RAF1: n = 10; COT: n = 10; MEKK2: n = 1; MLK1: n = 1) to yield a consensus expression vector. GSEA+Hallmark and ULM+CollecTRI were applied with identical parameters to the primary analysis. LINCS landmark gene coverage of the CollecTRI regulon was confirmed ≥5 targets for all focus TFs.

### Cross-dataset consistency analysis

Peterson RNA-seq and LINCS L1000 NES and ULM t-scores were pooled into a single ERK-versus-ERK+JNK comparison. Each MAP3K × dataset combination was treated as one observation. ERK group: RAF1×Peterson, COT×Peterson, RAF1×LINCS, COT×LINCS (n = 4). ERK+JNK group: MLK1/MLK3/ZAK/TAK1/MEKK2/MEKK3×Peterson + MEKK2×LINCS + MLK1×LINCS (n = 8). Two-sided exact Wilcoxon rank-sum tests (coin package [25]) with Benjamini-Hochberg FDR were applied to GSEA NES values (three pre-specified Hallmarks) and ULM t-scores (four focus TFs) independently.

### Software and reproducibility

All analyses were performed in R (v4.4.2; R Core Team, 2024). Key packages: DESeq2 [19], apeglm [20], fgsea [21], decoupleR [6], OmnipathR [23], cmapR [24], coin [25], ggplot2 [26], ggbeeswarm [27], ggrepel [28], pheatmap [29]. Complete session information is deposited with the analysis scripts. All five analysis scripts are self-contained and self-installing. Analysis code and processed data tables are deposited at [GitHub/Zenodo — to be completed before submission] under CC-BY 4.0.

## Acknowledgements

We thank Amy F. Peterson, Kayla Ingram, and Sergi Regot (Johns Hopkins University) for depositing the RNA-seq data publicly under GEO accession GSE213882, without which this re-analysis would not have been possible. We also acknowledge the LINCS Consortium and the Broad Institute for generating and publicly releasing the LINCS L1000 Phase I dataset (GEO: GSE92742), which enabled independent cross-platform validation of the findings reported here.

## Author Contributions

M.S.: Formal analysis, Writing – original draft, Visualization. B.C.E.: Investigation, Writing – review & editing. R.K.d.P.: Investigation, Writing – review & editing. C.P.M.: Formal analysis, Writing – review & editing. N.T.: Conceptualization, Methodology, Writing – review & editing, Validation, Supervision, Funding acquisition. A.G.: Conceptualization, Supervision, Funding acquisition, Writing – review & editing, Project administration.

## Data Availability

RNA-seq data re-analysed here are available from GEO under accession GSE213882 (Peterson et al. [5]). LINCS L1000 Phase I data are available under GEO accession GSE92742 (Subramanian et al. [9]).

## Supplementary Figure Legends

**Supplementary Fig. S1.**
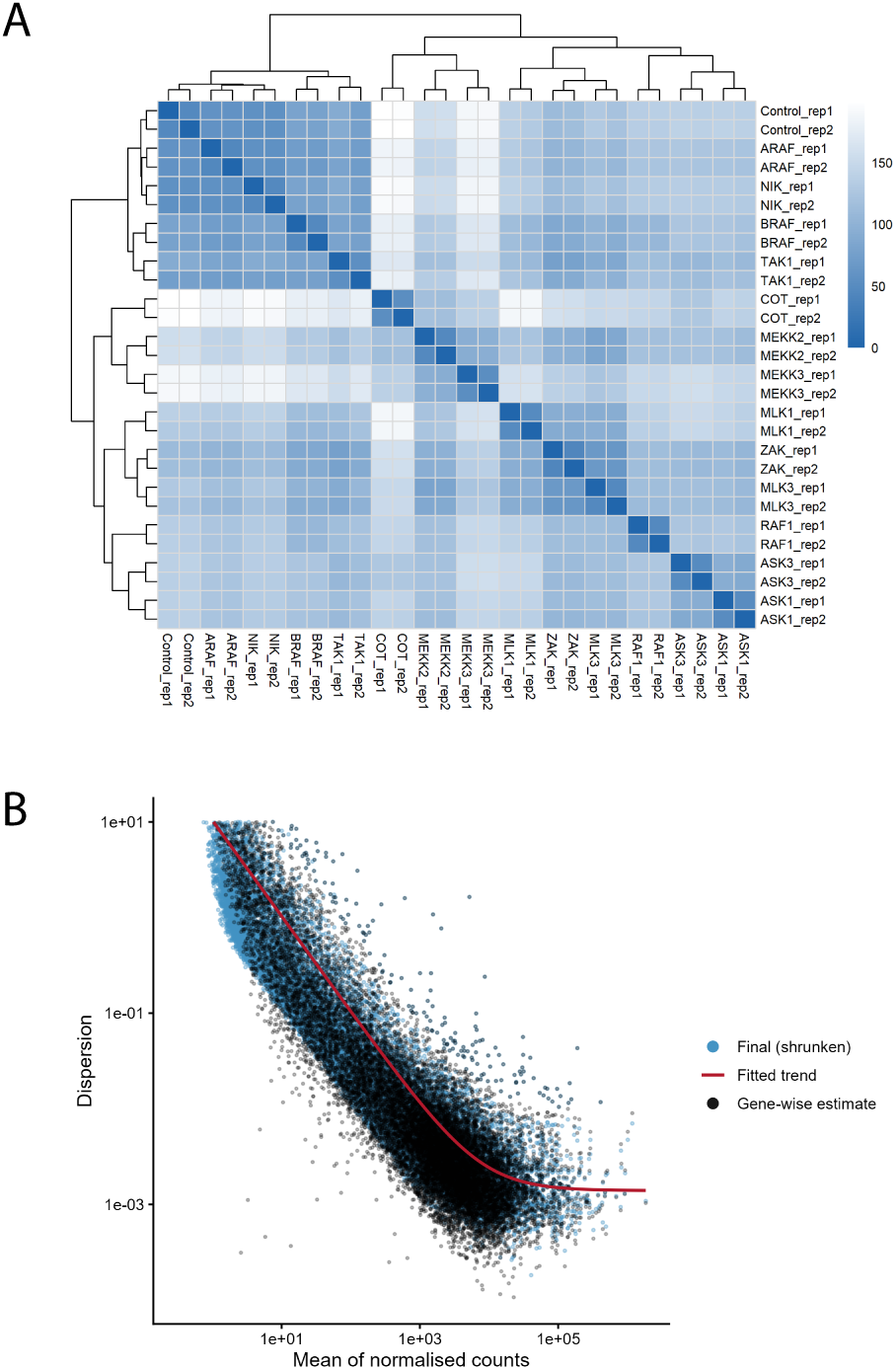
RNA-seq quality control. (a) Sample distance heatmap. Euclidean distances between all samples based on VST-normalised counts. Hierarchical clustering (Ward’s method). (b) DESeq2 dispersion-mean plot. Gene-wise dispersion estimates (grey) and fitted trend (red) across all MAP3K comparisons. Final shrunken dispersions in black.

**Supplementary Fig. S2.**
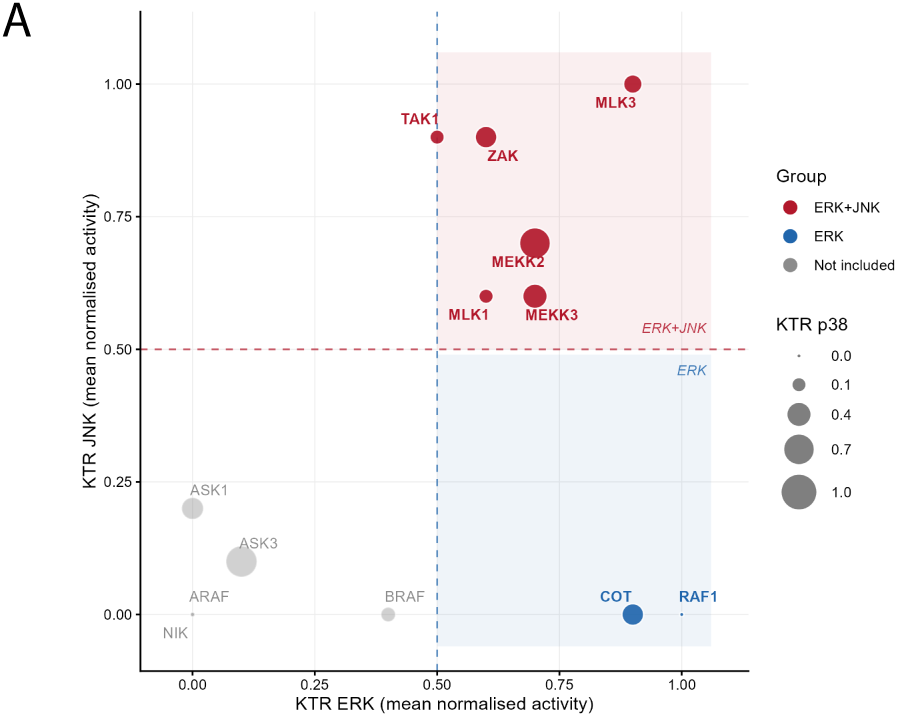
MAP3K group assignment based on KTR activity values. KTR activity values for ERK, JNK, and p38 across all MAP3Ks, based on data from Peterson et al. [5]. Dashed lines indicate thresholds used for group assignment. MAP3Ks excluded from primary analyses are shown in grey.

**Supplementary Fig. S3.**
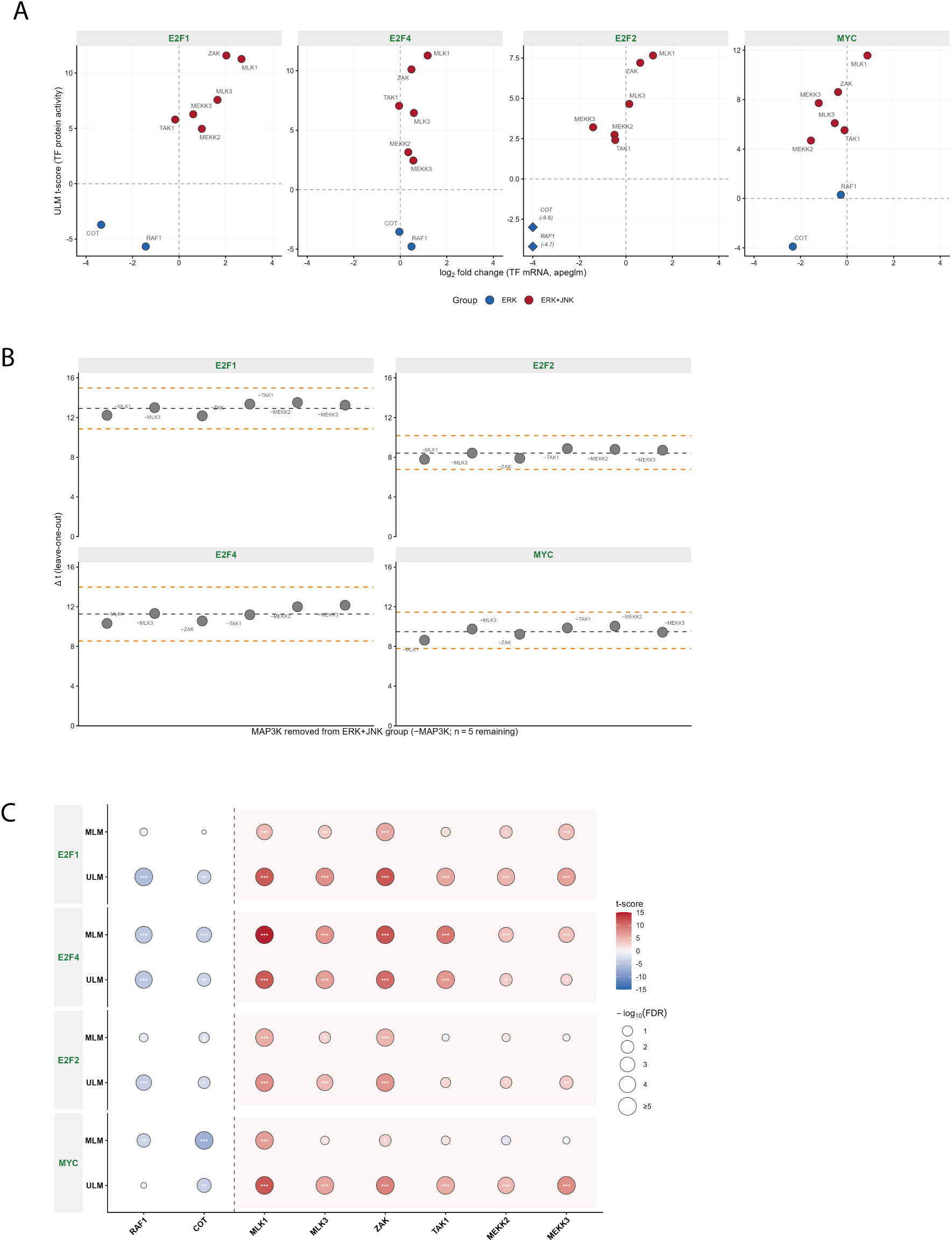
Method validation, sensitivity analysis, and TF–mRNA decoupling. (a) TF activity versus TF mRNA abundance. Scatter plot of ULM t-score (y-axis) versus TF mRNA log₂ fold-change from DESeq2 apeglm (x-axis) for all four focus TFs across all eight MAP3Ks. Each point represents one MAP3K; colour indicates group (blue = ERK, red = ERK+JNK). (b) Leave-one-out (LOO) sensitivity analysis. Group mean t-score difference (Δt = mean ERK+JNK − mean ERK) for each focus TF when each ERK+JNK MAP3K is removed in turn (n = 5 remaining). Each point represents one MAP3K removal. Grey dashed line: Δt from the full ERK+JNK group (n = 6). Orange dashed lines: 95% bootstrap confidence interval of the full-group Δt. (c) Method concordance between CollecTRI+ULM and DoRothEA+MLM. Interleaved dot plot of TF activity t-scores for four focus TFs across all MAP3Ks. Each panel shows ULM (CollecTRI; top) and MLM (DoRothEA A+B; bottom). Pearson correlation across all TF × MAP3K pairs (n = 32): r = 0.77 (p < 0.001).

**Supplementary Fig. S4.**
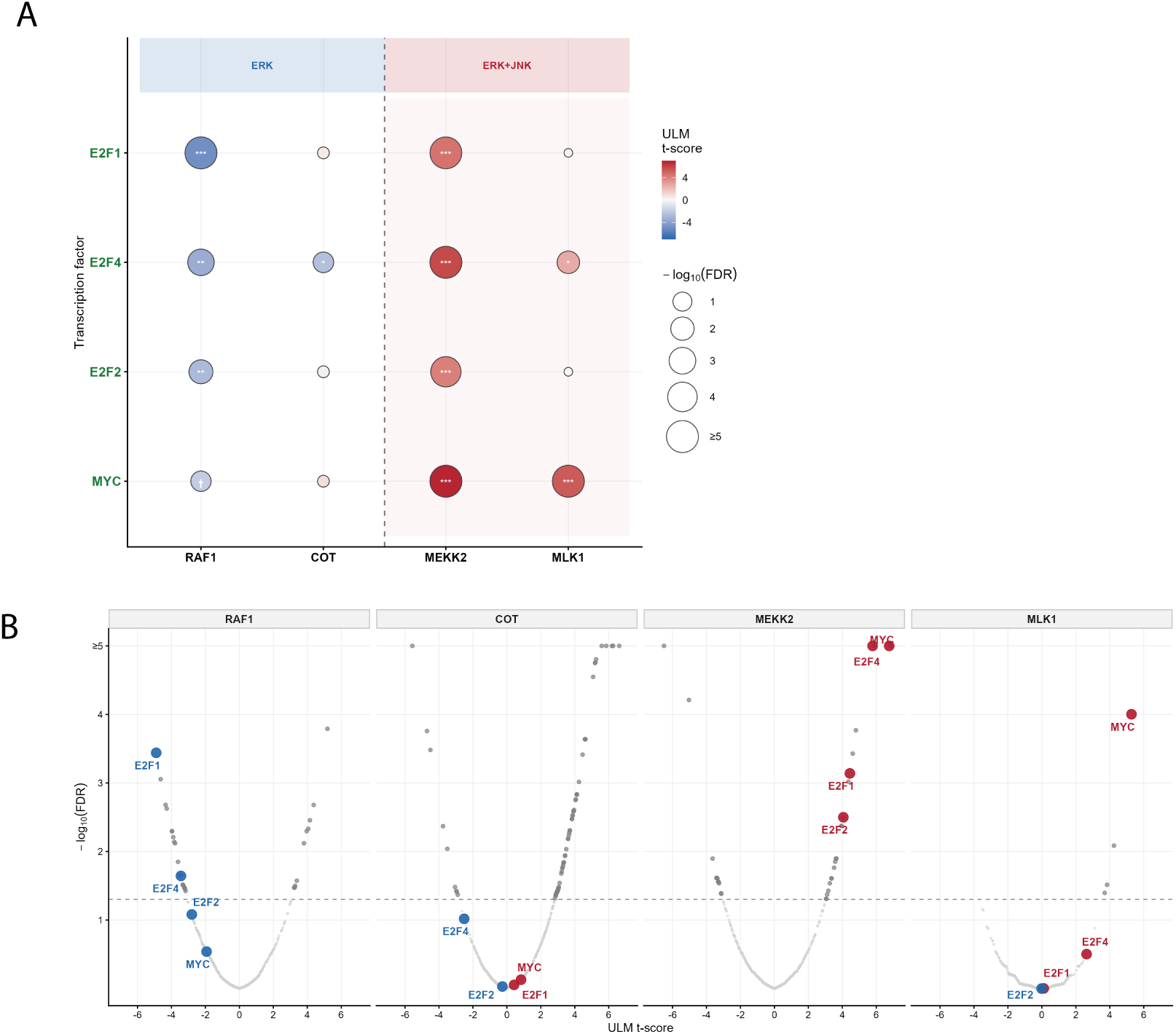
LINCS L1000 external validation. (a) ULM dot plot — LINCS L1000. ULM+CollecTRI t-scores for the four pre-specified focus TFs across all four LINCS MAP3K signatures. Colour and size encoding as in Fig. 3a. (b) Volcano plot across all CollecTRI TFs — LINCS L1000. All CollecTRI TFs tested by ULM on the LINCS consensus matrix. Panels faceted by MAP3K (RAF1, COT, MEKK2, MLK1). Encoding as in Fig. 3b.

